# An e-cigarette aerosol generation, animal exposure and toxicants quantification system to characterize *in vivo* nicotine kinetics in arterial and venous blood

**DOI:** 10.1101/2022.10.12.511981

**Authors:** Jushan Zhang, Mo Xue, Rong Pan, Yujie Zhu, Zhongyang Zhang, Haoxiang Cheng, Johan L M Björkegren, Jia Chen, Zhiqiang Shi, Ke Hao

**Affiliations:** Department of Respiratory Medicine, Shanghai Tenth People’s Hospital, Tongji University, Shanghai, China; College of Environmental Science and Engineering, Tongji University, Shanghai, China; Smoore Research Institute, Smoore International, Shenzhen, China; Department of Genetics and Genomic Sciences, Icahn School of Medicine at Mount Sinai, New York, NY, USA; Integrated Cardio Metabolic Centre, Department of Medicine, Karolinska Institutet, Karolinska Universitetssjukhuset, Huddinge, Sweden; Department of Environmental Medicine and Public Health, Icahn School of Medicine at Mount Sinai, New York, NY, USA

**Author notes:** These authors contribute equally (Jushan Zhang, Mo Xue).

## Abstract

The increasing e-cigarette use worldwide presents an urgent need to characterize their nicotine delivery property, brain stimulation and potential long-term health effects. We constructed an end-to-end system enabling combustible-cigarette (c-cigarette) and e-cigarette aerosol generation, animal exposure, and effect assessment. The system consists of (1) a 10-channel aerosol generator resembling human smoking/vaping scenarios, (2) nose-only and whole-body exposure chambers suitable for long- or short-duration studies, (3) a lab protocol for animal exposure and collecting arterial and venous blood <1 minute after the exposure, and (4) chromatograph and mass spectrometry to quantify nicotine concentrations in aerosol and biospecimens. We applied the system in a proof-of-principle study characterizing *in vivo* nicotine delivery after e-cigarette aerosol inhalation. Groups of Sprague-Dawley rats were exposed to e-cigarette aerosols for 1, 2 and 4 minutes, respectively. Arterial and venous blood samples were collected immediately after the exposure. We also directly compared nose-only and whole-body exposure approaches. After nose-only e-cigarette aerosol exposure, the nicotine concentration in arterial blood was substantially higher (11.32 ng/mL in average) than in veins. Similar arterio-venous concentration difference was observed in whole-body exposure experiments. In summary, we described a complete system ideal for e- and c-cigarette *in vivo* nicotine kinetics and long-term health research. Our findings highlight arterial blood as the suitable bio-specimen for e-cigarette nicotine delivery studies.

**Highlight:**

- We constructed a combustible- and e-cigarette aerosol generation - exposure - effect assessment system resembling real world human smoking/vaping scenarios.
- Proof-of-principle study characterized *in vivo* nicotine delivery from e-cigarette aerosol to arterial and venous blood at high temporal resolution.
- After exposure, the nicotine concentration was substantially higher (11.32 ng/mL) in arterial blood than in veins.
- Our results suggest arterial blood as the suitable bio-specimen to study nicotine delivery and brain stimulation.

## Introduction

Electronic cigarettes (e-cigs) are non-combustible products that generate inhalable aerosols from e-liquid^1–3^. In 2018, 14.9% of adults in the US had ever used an e-cig, and 3.2% were current e-cig users^1^. Rates of e-cig use (*i.e*., vaping) are highest in young people^2, 3^. Among 8th, 10th, and 12th grade students in the US, prevalence of e-cig use in 2018 was 9.7%, 20%, and 25%, respectively^4, 5^. Multiple types of e-cigs are currently on the market, such as e-cigs with and without nicotine. Besides nicotine, e-cig aerosols contain a variety of toxicants, some also found in combustible-cigarettes (c-cigs), *e.g*., reactive oxygen species [ROS] and metals,^6, 7^ as well as toxicants specific to e-cigs, *e.g*., carbonyls released from heating e-liquid solvents (propylene glycol [PG] and vegetable glycerin [VG])^3,8^. There are increasing concerns on the acute and chronic health hazard attributable to e-cig use^8, 9^. E-cigarette or Vaping Use-Associated Lung Injury (EVALI) is a severe lung illness^10, 11^. As of February 18, 2020, the US Centers for Disease Control (CDC) reported 2,807 EVALI cases hospitalized^10, 11^, among which 86% reported any use of THC (Tetrahydrocannabinol)-containing products, 64% reported any use of nicotine-containing products, 52% reported any use of both THC-containing products and nicotine-containing products, 34% reported exclusive use of THC-containing products, and 11% reported exclusive use of nicotine-containing products^10^. Further, as of February 18, 2020, US CDC reported 68 confirmed EVALI deaths^10, 11^. Besides acute injury and death, epidemiological studies indicated e-cig might increase risk of a number of chronic illness including chronic obstructive pulmonary disease (COPD), neurodegenerative disease, and cardiovascular disease^12, 13^.

Complementary to human observational studies, animal models are important tools to investigate e-cigs’ health effects. Animal exposure studies can be conducted under well controlled conditions and are able to dissect etiological mechanisms at organ, tissue, cellular and molecular levels. A variety of e-cig aerosol generation and animal exposure systems have been invented and some are now commercially available^14–16^. The key functional subunits in e-cig aerosol generators are e-cig changing robots and/or e-cig puffing triggers, aerosol pumping systems, inhalation exposure chambers and gas monitoring peripherals^17^. Specifically, there are two types of exposure chambers: whole-body and nose-only^18–21^. The nose-only system delivers aerosols primarily to the nasal regions, similar to human vaping situations; however, it requires the animal to be tightly restrained in a holding tube and only suitable for short duration exposures. In whole body exposure system, the animals are immersed in the test atmosphere and feasible for long duration exposure experiments.

The particulate matters (PM) from c-cig emission have relatively long lifespan (approximately 1.4 hours)^22^, while e-cig aerosol particles are liquid droplets in gas phase and of much shorter lifespan (*e.g*., 10-20 seconds)^17, 22^. Realtime assay found e-cigs aerosols have high mass concentration reaching >3000 μg/m^3^, however, the particles volatilized rapidly due to the high volatility of PG and VG^22^. Further, many e-cig aerosol components, including nicotine, have high boiling points and lower vapor pressures and tend to condense rapidly, especially when in contact with surfaces colder than the vapor^22, 23^. In nose-only exposure, the time from e-cig aerosol production to inhalation is less than one minute. In whole-body exposure experiments, the time from aerosol production to inhalation could be a few minutes, during which the aerosols undergo profound changes. It is possible that the animals inhale e-cig aerosols of different size and concentrations in nose-only and whole-body exposure settings, even the aerosols were generated in the same way. Multiple *in vivo* studies directly compared the nose-only and whole-body exposure of c-cig aerosols^19^ or metal PMs^18^, but not e-cig aerosols.

E-cig health studies typically employ chronic exposure strategy and measure the toxicant levels (*e.g*. nicotine) in venous blood. Venous nicotine levels represent the average effluent concentrations from all organs, and do not directly indicate nicotine levels in brain. Nicotine is lipid soluble and readily crosses the blood-brain barrier (BBB), therefore, nicotine rapidly reaches equilibrium between arterial blood and the brain. Post-inhalation nicotine level in the brain should be more accurately reflected in arteries than in veins. Studies showed, shortly after c-cig smoking, levels of nicotine in arterial plasma were more than double comparing to venous plasma^24, 25^, and the arterial nicotine concentration directly related to satisfaction and addiction of the c-cigarette products^24^. However, the arterial nicotine concentration or arterio-venous difference in e-cig users are not fully studied.

Aiming to fill the above knowledge gaps, we established an end-to-end system: e- and c-cig aerosol generation → animal exposure → *in vivo* toxicant assessment. The system includes four key elements. (1) Aerosol generation, a programmable multi-channel c- and e-cigarettes aerosol generator, which resembles human smoking behavior. (2) Nose-only and whole-body exposure chambers. (3) A lab protocol for animal exposure, surgery and arterial and venous blood collection <1 minute after the exposure. (4) Chemical assessment, gas chromatography and mass spectrometry for toxicant quantification in aerosol and bio-specimen. We comprehensively assessed the performance of this system, afterward, conducted a proof-of-principle study characterizing *in vivo* nicotine delivered in arterial and venous blood after e-cig aerosol inhalation.

## Materials and Methods

### Aerosol generation

The e- and c-cig aerosol generator consists of 10 independent channels controlled by programmable pumps (Fig 1A and 1B). A channel holds one e- or c-cig and operates in cycles. Each cycle lasts for 30 seconds, where the pump puffs for 3 seconds and produces 55mL cigarette aerosol, followed by 27 seconds of resting. Such maneuver mimics human smoking behavior, and the resting period allows the ecig atomizer to cool down between puffs. The 10 channels are programed to operate with a phase shift of 3 seconds, as the results, the system outputs aerosol stably at 1.1 L/min (Fig 1C). Zero air is introduced through an independent pathway to adjust the total aerosol volume and accommodate a variety of exposure chambers. In the aerosol generator, the cigarettes holders and all pipes were designed for easy removal and daily cleaning.

**Fig 1:**
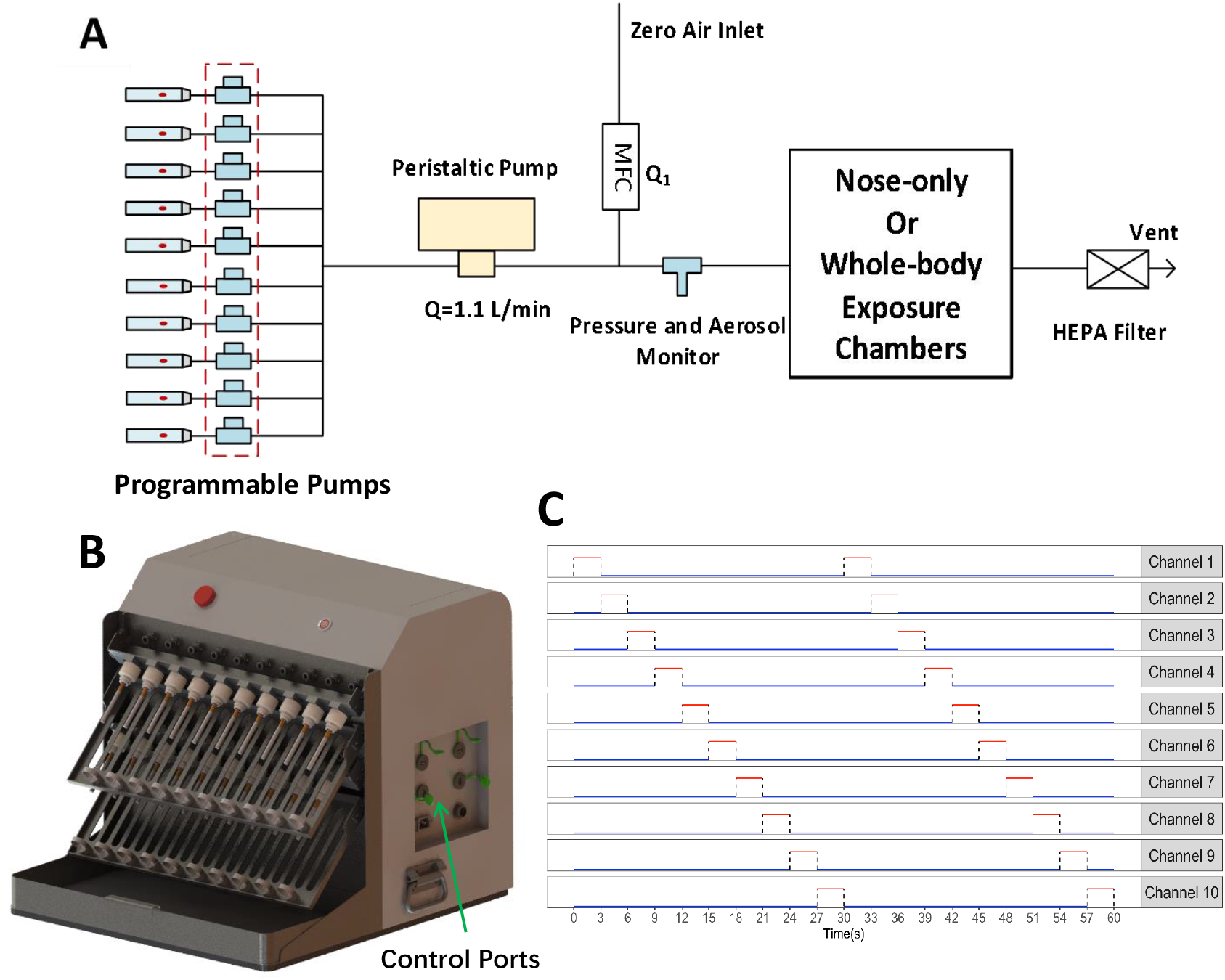
Aerosol generation systems. **A**. Diagram of system design. MFC, mass flow controller. **B**. Multichannel e- and c-cigarette aerosol generator. **C**. Operation cycles of the 10 independently programmable channels for 60 seconds. Red line, the channel is in cigarettes puff mode, and blue line, the channel is in resting/cooling mode.

In this study, the zero air flow rate was set at 8.9 L/min, and the total output of the cigarette aerosol generator was 10L/min. In the c-cig exposure experiments, we used reference cigarettes 1R6F (Kentucky Tobacco Research & Development Center, Lexington, KY, USA). In e-cig exposure experiments, the eliquid was composed of 50% VG, 40% propylene glycol PG, 5% nicotine, and 5% benzoic acid. The eliquid was heated by porous ceramic coils (Smoore International, Shenzhen, China).

### Quantification of nicotine concentration in e- and c-cigarette aerosols

To avoid cross-contamination, the e- and c-cig aerosol experiments were performed separately following the same protocol. First, we produced e- or c-cig aerosols using the aerosol generator. Second, the aerosols were collected by 44 mm Cambridge filter pad (Whatman Lab), then eluted by isopropanol solution. Third, the nicotine concentration was quantified by gas chromatography (Agilent 7890B) equipped with a flame ionization detector (FID). We utilized the Aglient DB-WAX UI columns (30m*0.25mm*0.25um) following manufacturer recommended protocols. A calibration curve was established (Fig 2) using solutions of known nicotine concentrations, where we observed high linearity indicating the experiments operated within the dynamic range of excellent reliability and accuracy. Finally, we quantified the nicotine concentrations in e-cig or c-cig aerosols based on the calibration curve (Fig 2).

**Fig 2.**
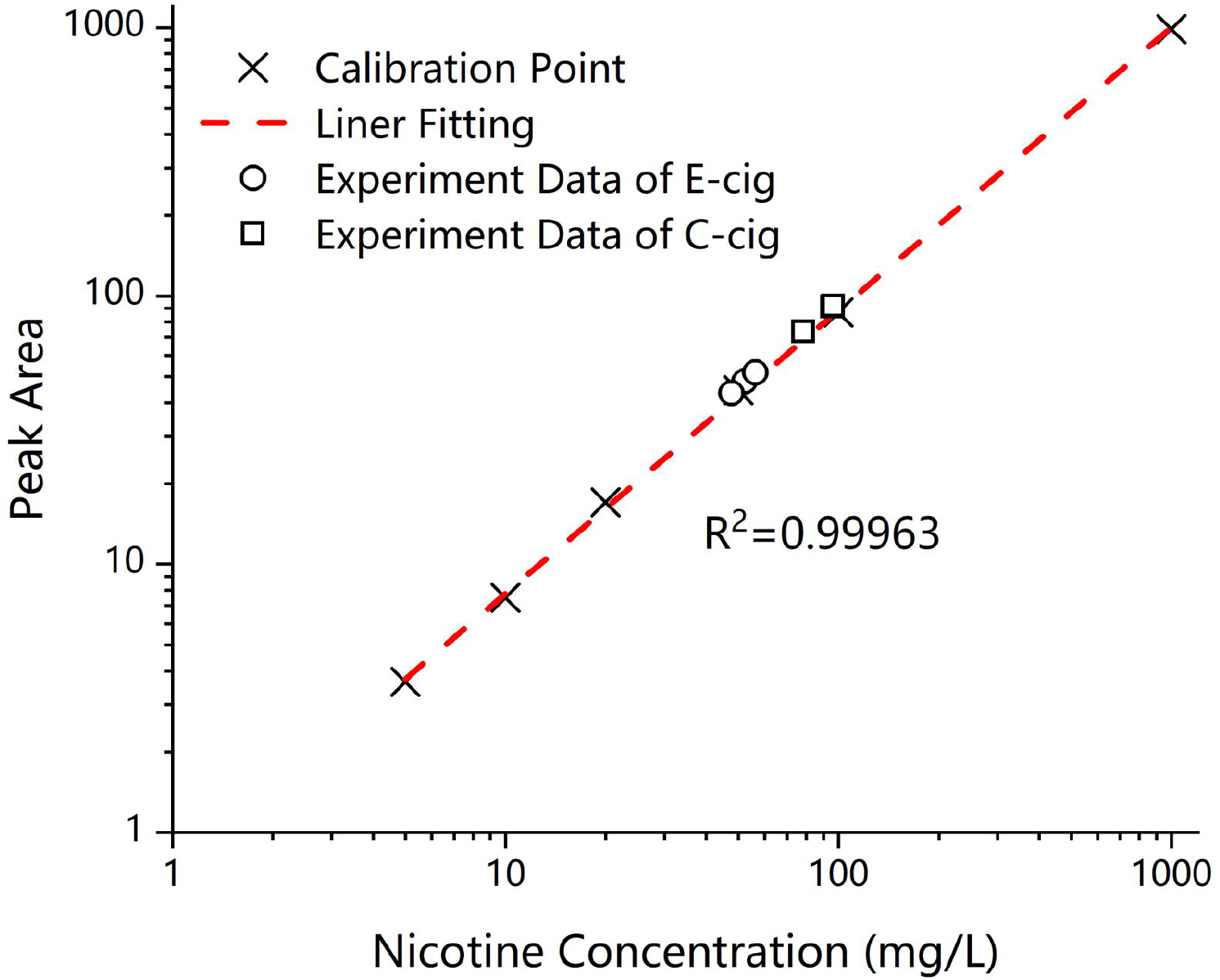
Calibration curve in quantifying the e-cigarette aerosol nicotine concentrations. Peak area, area under the curve.

### Nose-only and whole-body exposure systems

We employed commercially available nose-only and whole-body exposure chambers (TOW-INT TECH, Shanghai, China). The nose-only exposure manifold consisted of twelve ports (Fig 3A), connecting to restraining tubes of different size that accommodate various animal model (*e.g*., mouse and rat) or development stages (young vs. adult). Rotating fittings allows for fast loading and retrieving the animals. The aerosols enter an inlet plenum that directs the flow to each restraining tube. All restraining tubes receive homogenous aerosol with < ±5% concentration variation. We operate the exposure manifold inside a fume hood. The exhaust air (exhaled and excess aerosol flow) flows through high-efficiency particulate air (HEPA) filter and then release to the fume hood’ exhaust duct^14, 26^.

**Fig 3:**
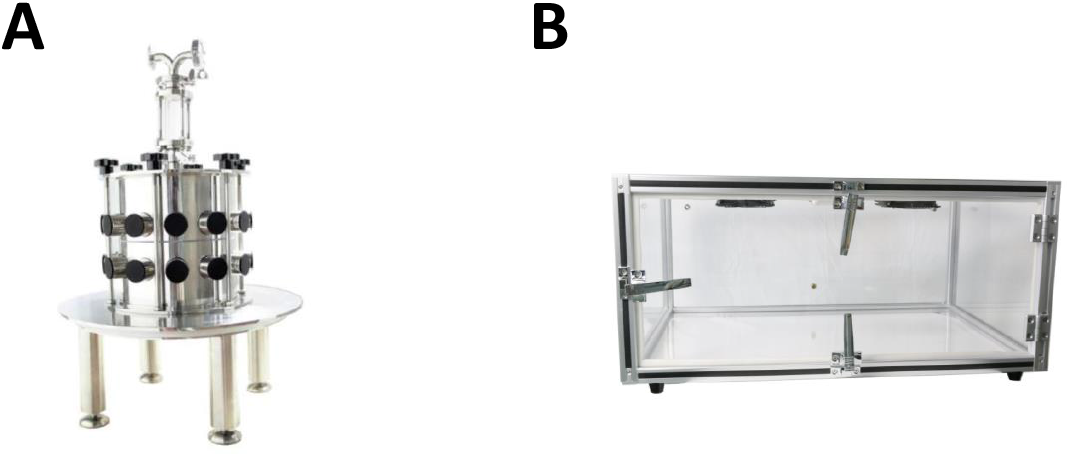
Nose-only (**A**) and whole-body (**B**) exposure chambers.

The whole-body exposure chamber is suitable for long-duration exposure studies. The dimension of the cuboid acrylic chamber was 788mm X 358mm X 543mm (W X H X D) with side hinged opening (Fig 3B). The chamber houses up to four mice cages or two rat cages. The side hinged opening allowed rapid handling of the animal cage. This feature is critical for nicotine kinetics studies, since animals need to be quickly retrieved from the exposure chamber for arterial and veinous blood collection. Aerosol enters the chamber from the injection ports located on both right and left sides. Two custom-built fans above the animal cage ensure uniform aerosol distribution. Humidity, temperature, O_2_, PM_2.5_ and CO levels inside the chamber are monitored by sensors in real-time. The exhaust filtration is same as in the nose-only exposure system. We operate whole-body exposure chamber inside a fume hood.

It should be noted that the e-/c-cig aerosol generator only connects to one exposure chamber (either nose-only or whole-body) in experiments. The generator’s output flow (10L/min) was not adequate to supply both chambers simultaneously.

### E-cigarette nicotine kinetics experiments: exposure and biospecimen collection

Eighteen male Sprague-Dawley rats of 6 weeks old and body weight of 180–200g were used in the exposure and nicotine kinetics study. The animals were kept in the vivarium under a 12-hr light/dark cycle and had *ad libitum* access to food and water. We equally divided the 18 rats into six groups. Three groups of mice were exposed to e-cig aerosols for 1 minute, 2 minutes, and 4 minutes, respectively, using nose-only method. Three groups of mice were exposed to e-cig aerosols for 1 minute, 2 minutes, and 4 minutes, respectively, using whole-body method (Fig 4). Nose-only and whole-body exposure experiments were conducted separately.

**Fig 4:**
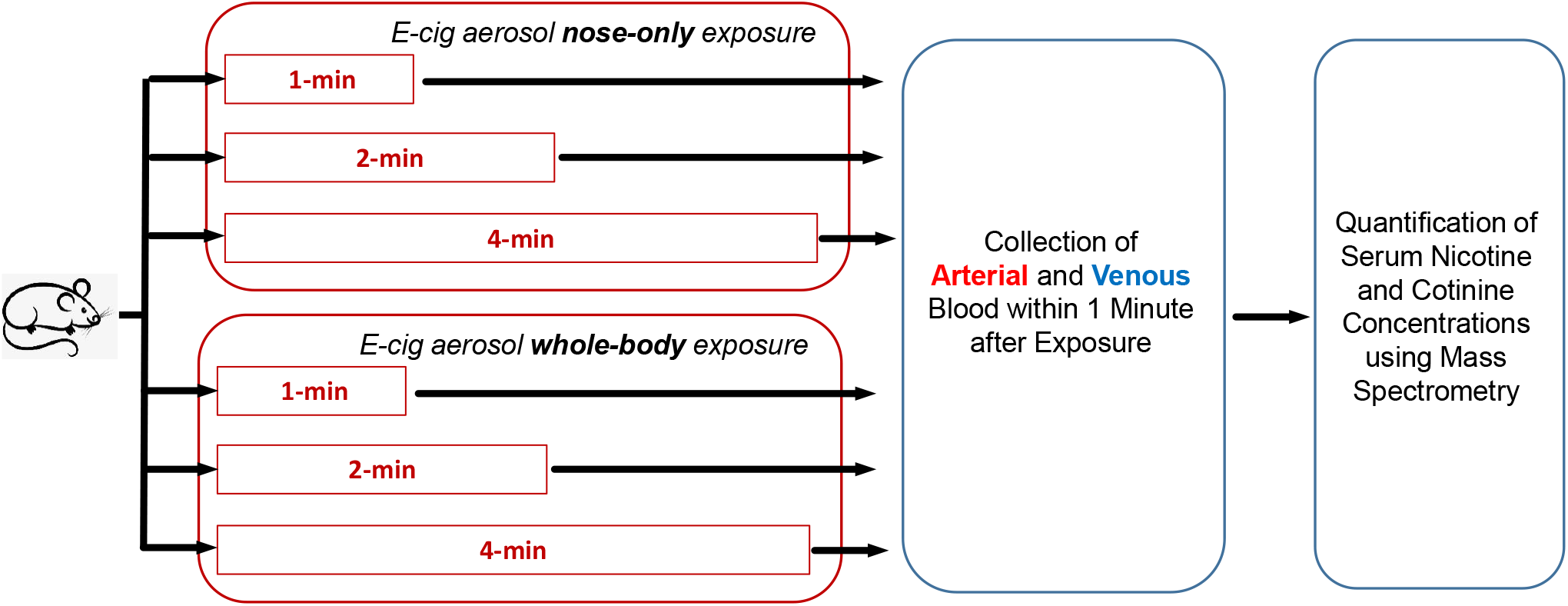
Design of the e-cig aerosol *in vivo* nicotine delivery study.

The nicotine kinetics experiment required precise control of exposure duration (*e.g*., 1 minute) and blood collection immediately after exposure. As a feasible strategy, we conducted the experiment using one animal at a time. In another word, only one rat was placed into the exposure chamber during each experiment. In detail, we firstly pumped e-cig aerosol into the exposure chamber for 5 minutes allowing the system reaching steady state. Second, we quickly placed one rat to the nose-only exposure manifold (nose-only method) or the exposure chamber (whole-body methods) to start the exposure. Third, the exposure lasted for designed duration (*e.g*., 1 minute). Fourth, the rat was quickly retrieved from the nose-only manifold or exposure chamber. Then, we immediately euthanized the rat and collected arterial blood (0.6mL) from abdominal aorta and venous blood (0.6mL) from inferior vena cava. The blood samples were centrifuged at 1,100g for 20 □ min at 4□°C to remove cells and clotting factors, afterward stored at −80°C.

We repeated the above steps one rat at a time, and completed the exposure and sample collection experiments on the eighteen rats.

### Quantification of nicotine and cotinine concentration in serum

200μL serum was pipetted into 1mL tube. 200μL internal standard (solution of nicotine-D4 and cotinine-D3) and 4% phosphoric acid solution were also added. After vortex mixing, the solutions went through MCX solid phase extraction plate, eluted by acetonitrile ammoniate, afterward, dried using a nitrogen evaporator. The dried sample was redissolved into 200μL mobile phase, followed by centrifuge at 5,000 rpm for 10 minutes. The supernatant was loaded into the sample bottle for UPLC-MS/MS analysis. The HPLC column (Aglient Poroshell 120 Hilic 2.7μm, 2.1×100mm) was maintained at 35°C, and 5μl sample was injected into the column. In liquid chromatography assay, the ion source and ion transfer line were all maintained at 350°C. Sheath flow, auxiliary flow and purge flow were 42 Bar, 10 Bar and 0 Bar, respectively.

Mass spectrometric detection was performed using Thermo Scientific Ultimate 3000-TSQ Quantiva Mass Spectrometer in positive ESI mode. The mass spectrometric parameters were optimized for both nicotine and cotinine and their respective internal standards to the following protonated ions: m/z 163 → 117 [M + H]^+^(nicotine), m/z 167→ 121.00 [M + H]^+^(nicotine-D4), m/z 177 → 80 [M + H]^+^(cotinine) and m/z 180→ 80[M + H]^+^(cotinine-D3) to reach maximum sensitivity. Calibration curves for nicotine and cotinine were established using sample of known concentrations of 0.1, 0.5, 1, 5, 10, 50, 100, 200 ng/mL. Shown in Fig 5, both calibration curves demonstrated high linearity, indicating the experiments operated in dynamic range of excellent reliability and accuracy. Finally, we quantified the nicotine and cotinine concentration in serum following the above protocol (Fig 5).

**Fig 5.**
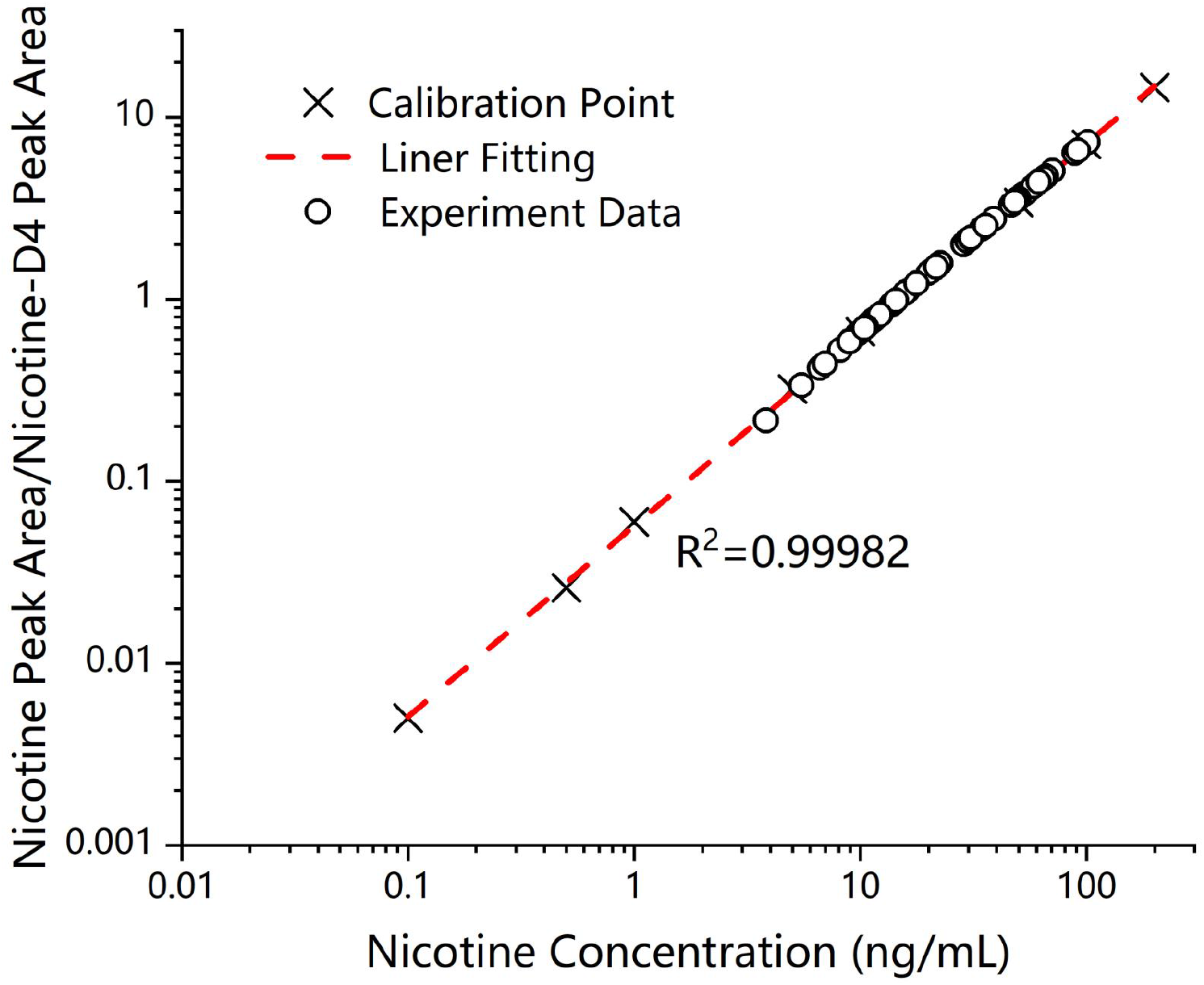
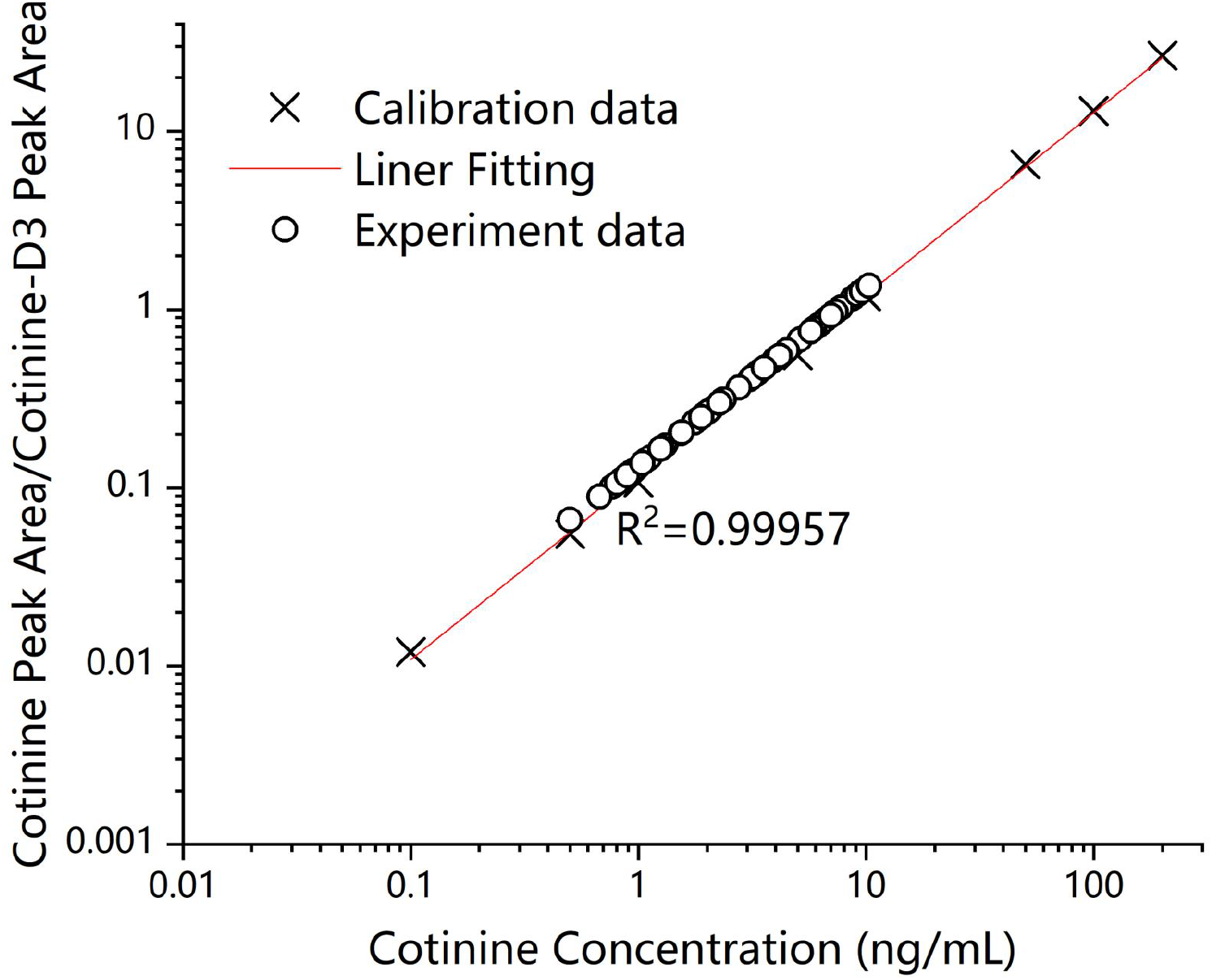
Calibration Curves in quantify the serum nicotine (**A**) and cotinine (**B**) concentrations. Peak area, area under the curve.

### Statistical analysis

Data analysis was conducted using R 4.1.1 statistical package. Means and standard deviations were presented for quantitative outcomes.

### System maintenance

We cleaned the system daily. (1) Animal hair or other debris was removed from the chamber and exhaust ducts. (2) The aerosol flow tubes were washed with water and detergent. To avoid potential cross-contamination between aerosols, we used dedicated sets of holders and tubes for e- and c-cigs.

## Results

### Cigarette aerosol generator performance and aerosol characterization

The e- and c-cigarette aerosol generator underwent extensively testing. The generator output aerosol stably at 1.1 L/min, and zero air was introduced through an independent pathway resulting in a steady flow of 10L/min into the exposure chamber. The nicotine concentration in the e- and c-cig aerosol entering the exposure chamber was quantified as 0.27 ± 0.01 mg/L and 0.42± 0.02 mg/L, respectively.

### Serum nicotine and cotinine concentration

Rats were exposed to e-cig aerosol by nose-only or whole-body methods for 1, 2, 4 minutes (Fig 4). Immediately after the exposure, we collected arterial and venous blood samples. The nicotine concentration in artery was substantially higher than that in vein. In nose-only exposure, the arterial blood nicotine concentration was 24.53 ng/mL, 37.67 ng/mL and 55.33 ng/mL after 1, 2 and 4 minutes of exposure, respectively (Fig 6 and Table S1). In contrast, the venous nicotine concentration was 12.97 ng/mL, 26.93 ng/mL and 43.67 ng/mL after exposure for 1, 2 and 4 minutes, respectively. On average, the arterial nicotine concentration was 11.32 ng/mL higher than venous concentration. Similar pattern was observed in the whole-body exposure experiments, where the arterial nicotine concentration was 12.82 ng/mL, 19.10 ng/mL and 38.23 ng/mL after 1, 2 and 4 minutes of exposure, respectively. In contrast, the venous nicotine concentration was 8.14 ng/mL, 15.03 ng/mL and 24.13 ng/mL after 1, 2 and 4 minutes of exposure, respectively. The two exposure methods also showed substantial nicotine delivery rate, where the nose-only exposure produced in average 15.79 ng/mL higher arterial blood nicotine concentration than whole-body method. Cotinine is the predominant metabolite of nicotine. As expected, cotinine had comparable arterial and venous concentrations in both nose-only and whole-body exposure experiments (Fig 7). We also observed higher blood cotinine levels among mice exposed to ecig aerosol by nose-only method than whole-body method.

**Fig 6:**
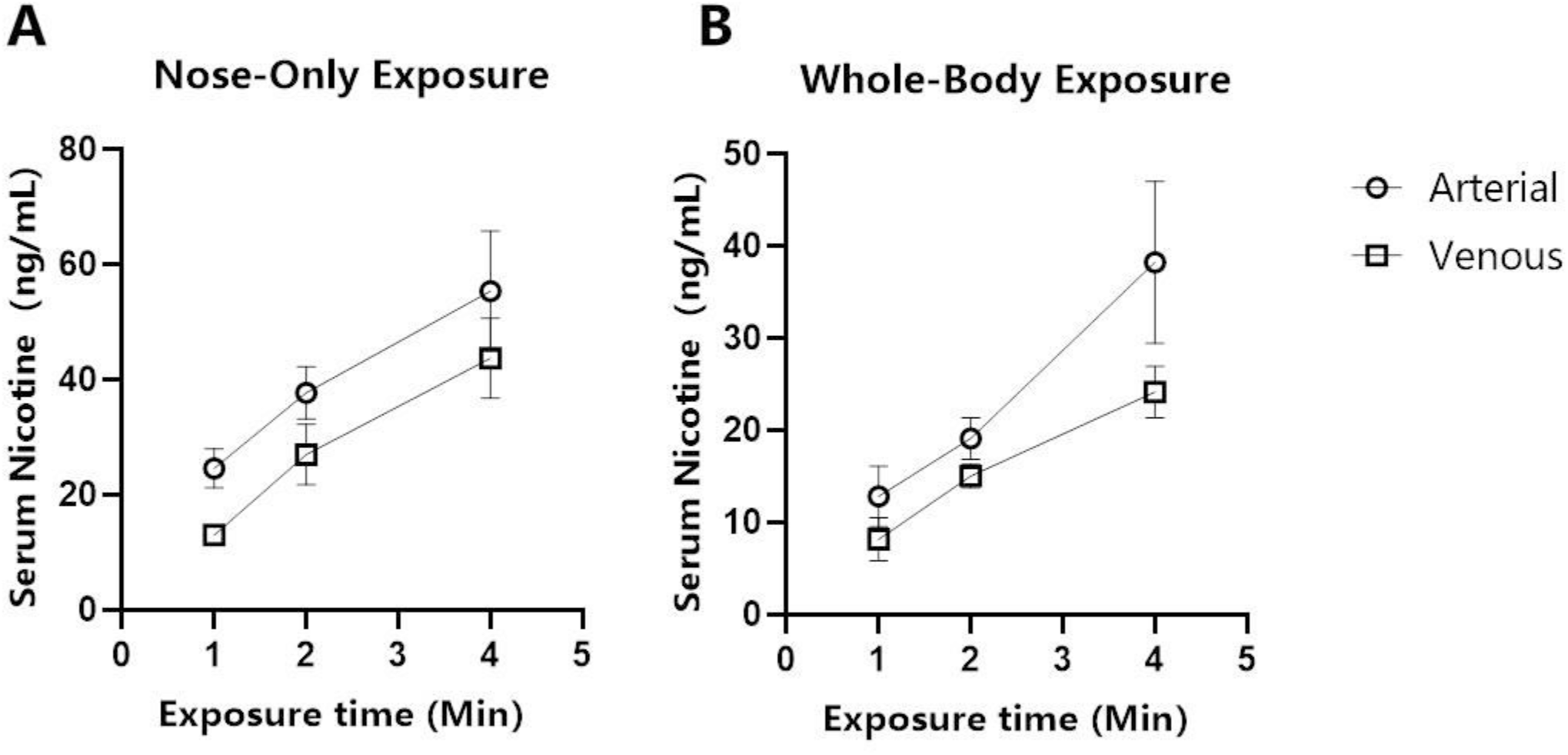
Blood nicotine level after e-cig aerosol exposure using nose-only (**A**) and whole-body (**B**) methods.

**Fig 7:**
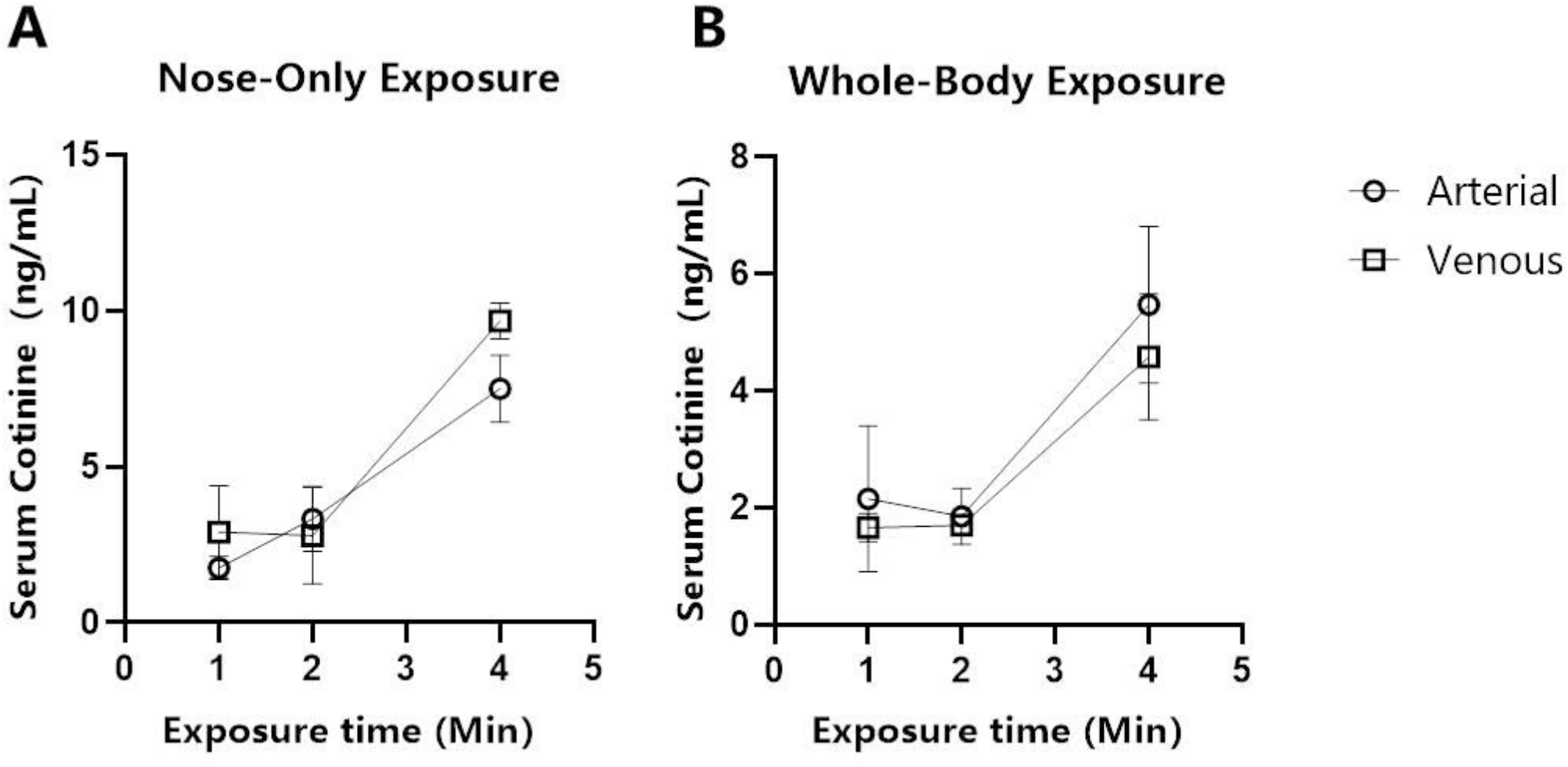
Blood cotinine level after e-cig aerosol exposure using nose-only (**A**) and whole-body (**B**) methods.

## Discussion

We describe an end-to-end system consisted of e- and c-cig aerosol generation, animal exposure, bio-specimen collection, and *in vivo* nicotine delivery assessment. The 10-channel aerosol generator resembles human smoking behavior and produces e- and c-cig aerosol at well-controlled rate and concentration. We conducted a proof-of-principle study and made the following key observations. (1) Consistent to previous reports^27^, the c-cig aerosols had higher nicotine concentration than aerosols of ecigs (5% nicotine e-liquid) (Fig 2). (2) The arterial blood nicotine concentration was substantially higher (11.32 ng/mL on average) than that in veins. We characterized the arterial blood nicotine boost (rate of nicotine concentration increase) after 1, 2 and 4 minutes of e-cig aerosol inhalation (Fig 6), which reflected the rate of nicotine delivered to the brain. (3) The nose-only and whole-body methods were suitable for short- and long-duration exposure, respectively, and nose-only method consistently produced higher arterial blood nicotine concentration than whole-body method.

The amount and rate of nicotine delivered to arterial blood are key factors determining user satisfaction and effectiveness of e-cigs in smoking cessation. Different types of e-cigs vary greatly in nicotine delivery properties^27^, and the e-liquid formulation is also important^28^. Randomized trial and cohort study showed e-cigs of low nicotine deliver rate failed to achieve impressive smoking cessation results^29, 30^. Unfortunately, existing e-cig nicotine delivery studies were mostly based on venous blood. Herein, we demonstrated the arterial blood nicotine concentrations were substantially higher than venous blood after e-cigarette use. Further, nicotine is lipid soluble. It easily crosses the BBB and quickly reaches equilibrium between arterial blood and brain tissues. Therefore, we argue arterial blood is the suitable bio-specimen to characterize e-cig nicotine delivery rate, brain stimulation, user satisfaction and self-perceived addiction.

In our system, it took less than 20 seconds for the aerosol to travel from e-/c-cigs to the exposure chamber. Given certain toxicants (*e.g*., ROS) had short half-life, minimizing the aerosol travel time was essential to resemble real-world vaping scenarios. We observed the nose-only exposure produced higher arterial nicotine concentration than whole-body method. In nose-only exposure, it only took <20 seconds from aerosol generation to inhalation. In whole-body exposure, aerosols could stay for minutes in the chamber (Fig 3B) before inhaled by animals, allowing nicotine to condense given its high boiling points and lower vapor pressures. We designed the lab protocol feasible for nicotine kinetics study at high temporal resolution. Only one rat was employed in each experiment enabling precise control of exposure duration (*e.g*., 1 minute) and blood collection immediately after exposure.

In summary, we constructed an e- and c-cig aerosol generation – exposure – effect assessment system resembling real world smoking/vaping. The proof-of-principle study characterized *in vivo* nicotine delivery at high temporal resolution after e-cig use. We particularly emphasize the importance of arterial blood in nicotine delivery and kinetics studies.

## Reference

1. Creamer, M. R.; Wang, T. W.; Babb, S.; Cullen, K. A.; Day, H.; Willis, G.; Jamal, A.; Neff, L., Tobacco Product Use and Cessation Indicators Among Adults - United States, 2018. MMWR Morb Mortal Wkly Rep 2019, 68, (45), 1013–1019.

2. Gottlieb, M. A., Regulation of E-Cigarettes in the United States and Its Role in a Youth Epidemic. Children (Bosel) 2019, 6, (3).

3. Gotts, J. E.; Jordt, S.-E.; McConnell, R.; Tarran, R., What are the respiratory effects of e-cigarettes? BMJ 2019, 366, 15275.

4. Patrick, M. E.; O’Malley, P. M.; Kloska, D. D.; Schulenberg, J. E.; Johnston, L. D.; Miech, R. A.; Bachman, J. G., Novel psychoactive substance use by US adolescents: Characteristics associated with use of synthetic cannabinoids and synthetic cathinones. Drug Alcohol Rev 2016, 35, (5), 586–90.

5. Miech, R.; Johnston, L.; O’Malley, P. M.; Bachman, J. G.; Patrick, M. E., Adolescent Vaping and Nicotine Use in 2017-2018 - U.S. National Estimates. N Engl J Med 2019, 380, (2), 192–193.

6. Haddad, C.; Salman, R.; El-Hellani, A.; Talih, S.; Shihadeh, A.; Saliba, N. A., Reactive Oxygen Species Emissions from Supra- and Sub-Ohm Electronic Cigarettes. J Anal Toxicol 2019, 43, (1), 45–50.

7. Zhao, J.; Zhang, Y.; Sisler, J. D.; Shaffer, J.; Leonard, S. S.; Morris, A. M.; Qian, Y.; Bello, D.; Demokritou, P., Assessment of reactive oxygen species generated by electronic cigarettes using acellular and cellular approaches. Journal of hazardous materials 2018, 344, 549–557.

8. Olmedo, P.; Goessler, W.; Tanda, S.; Grau-Perez, M.; Jarmul, S.; Aherrera, A.; Chen, R.; Hilpert, M.; Cohen, J. E.; Navas-Acien, A.; Rule, A. M., Metal Concentrations in e-Cigarette Liquid and Aerosol Samples: The Contribution of Metallic Coils. Environ Health Perspect 2018, 126, (2), 027010–027010.

9. Glantz, S. A.; Bareham, D. W., E-Cigarettes: Use, Effects on Smoking, Risks, and Policy Implications. Annu Rev Public Health 2018, 39, 215–235.

10. https://www.cdc.gov/media/releases/2019/p1028-first-analysis-lung-iniury-deaths.html.

11. https://www.cdc.gov/tobacco/basic_information/e-cigarettes/severe-lung-disease.html.

12. Merecz-Sadowska, A.; Sitarek, P.; Zielinska-Blizniewska, H.; Malinowska, K.; Zajdel, K.; Zakonnik, L.; Zajdel, R., A Summary of In Vitro and In Vivo Studies Evaluating the Impact of E-Cigarette Exposure on Living Organisms and the Environment. Int J Mol Sci 2020, 21, (2).

13. Lerner, C. A.; Sundar, I. K.; Yao, H.; Gerloff, J.; Ossip, D. J.; McIntosh, S.; Robinson, R.; Rahman, I., Vapors produced by electronic cigarettes and e-juices with flavorings induce toxicity, oxidative stress, and inflammatory response in lung epithelial cells and in mouse lung. PLoS One 2015, 10, (2), e0116732.

14. Wong, B. A., Inhalation exposure systems: design, methods and operation. Toxicol Pathol 2007, 35, (1), 3–14.

15. Li, X.; Nie, C.; Shang, P.; Xie, F.; Liu, H.; Xie, J., Evaluation method for the cytotoxicity of cigarette smoke by in vitro whole smoke exposure. Exp Toxicol Pathol 2014, 66, (1), 27–33.

16. Hilpert, M.; Ilievski, V.; Coady, M.; Andrade-Gutierrez, M.; Yan, B.; Chillrud, S. N.; Navas-Acien, A.; Kleiman, N. J., A custom-built low-cost chamber for exposing rodents to e-cigarette aerosol: practical considerations. Inhal Toxicol 2019, 31, (11-12), 399–408.

17. Zweier, J. L.; Shalaan, M. T.; Samouilov, A.; Saleh, I. G.; El-Mahdy, M. A., Whole body electronic cigarette exposure system for efficient evaluation of diverse inhalation conditions and products. Inhal Toxicol 2020, 32, (13-14), 477–486.

18. Oyabu, T.; Morimoto, Y.; Izumi, H.; Yoshiura, Y.; Tomonaga, T.; Lee, B.-W.; Okada, T.; Myojo, T.; Shimada, M.; Kubo, M.; Yamamoto, K.; Kawaguchi, K.; Sasaki, T., Comparison between whole-body inhalation and nose-only inhalation on the deposition and health effects of nanoparticles. Environ Health Prev Med 2016, 21, (1), 42–48.

19. Serré, J.; Tanjeko, A. T.; Mathyssen, C.; Vanherwegen, A.-S.; Heigl, T.; Janssen, R.; Verbeken, E.; Maes, K.; Vanaudenaerde, B.; Janssens, W.; Gayan-Ramirez, G., Enhanced lung inflammatory response in whole-body compared to nose-only cigarette smoke-exposed mice. Respiratory Research 2021, 22, (1), 86.

20. Kogel, U.; Wong, E. T.; Szostak, J.; Tan, W. T.; Lucci, F.; Leroy, P.; Titz, B.; Xiang, Y.; Low, T.; Wong, S. K.; Guedj, E.; Ivanov, N. V.; Schlage, W. K.; Peitsch, M. C.; Kuczaj, A.; Vanscheeuwijck, P.; Hoeng, J., Impact of whole-body versus nose-only inhalation exposure systems on systemic, respiratory, and cardiovascular endpoints in a 2-month cigarette smoke exposure study in the ApoE(-/-) mouse model. J Appl Toxicol 2021.

21. Cheng, Y. S.; Bowen, L.; Rando, R. J.; Postlethwait, E. M.; Squadrito, G. L.; Matalon, S., Exposing Animals to Oxidant Gases. Proceedings of the American Thoracic Society 2010, 7, (4), 264–268.

22. Lampos, S.; Kostenidou, E.; Farsalinos, K.; Zagoriti, Z.; Ntoukas, A.; Dalamarinis, K.; Savranakis, P.; Lagoumintzis, G.; Poulas, K., Real-Time Assessment of E-Cigarettes and Conventional Cigarettes Emissions: Aerosol Size Distributions, Mass and Number Concentrations. Toxics 2019, 7, (3).

23. Margham, J.; McAdam, K.; Forster, M.; Liu, C.; Wright, C.; Mariner, D.; Proctor, C., Chemical Composition of Aerosol from an E-Cigarette: A Quantitative Comparison with Cigarette Smoke. Chem Res Toxicol 2016, 29, (10), 1662–1678.

24. Henningfield, J. E.; Stapleton, J. M.; Benowitz, N. L.; Grayson, R. F.; London, E. D., Higher levels of nicotine in arterial than in venous blood after cigarette smoking. Drug Alcohol Depend 1993, 33, (1), 23–9.

25. Henningfield, J. E.; London, E. D.; Benowitz, N. L., Arterial-Venousn Differences in Plasma Concentrations of Nicotine After Cigarette Smoking. JAMA 1990, 263, (15), 2049–2050.

26. Buckley, A.; Warren, J.; Hodgson, A.; Marczylo, T.; Ignatyev, K.; Guo, C.; Smith, R., Slow lung clearance and limited translocation of four sizes of inhaled iridium nanoparticles. Part Fibre Toxicol 2017, 14, (1), 5.

27. Yingst, J. M.; Foulds, J.; Veldheer, S.; Hrabovsky, S.; Trushin, N.; Eissenberg, T. T.; Williams, J.; Richie, J. P.; Nichols, T. T.; Wilson, S. J.; Hobkirk, A. L., Nicotine absorption during electronic cigarette use among regular users. PLOS ONE 2019, 14, (7), e0220300.

28. Talih, S.; Balhas, Z.; Eissenberg, T.; Salman, R.; Karaoghlanian, N.; El Hellani, A.; Baalbaki, R.; Saliba, N.; Shihadeh, A., Effects of user puff topography, device voltage, and liquid nicotine concentration on electronic cigarette nicotine yield: measurements and model predictions. Nicotine Tob Res 2015, 17, (2), 150–7.

29. Bullen, C.; Howe, C.; Laugesen, M.; McRobbie, H.; Parag, V.; Williman, J.; Walker, N., Electronic cigarettes for smoking cessation: a randomised controlled trial. Lancet 2013, 382, (9905), 1629–37.

30. Kalkhoran, S.; Glantz, S. A., E-cigarettes and smoking cessation in real-world and clinical settings: a systematic review and meta-analysis. Lancet Respir Med 2016, 4, (2), 116–28.

