## Supplementary material for "An e-cigarette aerosol generation, animal exposure and toxicants quantification system to characterize *in vivo* nicotine kinetics in arterial and venous blood": Table S1

Table S1. Serum nicotine and cotinine concentration (mean ± SD) after nose-only or whole-body e-cigarettes aerosol exposure for 1,2,4 min

| **Exposure Method** |  | **Arterial Blood** | | **Venous Blood** | |
| --- | --- | --- | --- | --- | --- |
|  | **Exposure Time (min)** | **Serum Nicotine  （ng/mL）** | **Serum Cotinine  （ng/mL）** | **Serum Nicotine  （ng/mL）** | **Serum Cotinine  （ng/mL）** |
| **Nose-Only** | 1 | 24.53 ± 3.44 | 1.75 ± 0.38 | 12.97 ± 1.45 | 2.90 ± 1.49 |
|  | 2 | 37.67 ± 4.56 | 3.33 ± 1.04 | 26.93 ± 5.32 | 2.79 ± 1.55 |
|  | 4 | 55.33 ± 10.51 | 7.51 ± 1.08 | 43.67 ± 6.99 | 9.68 ± 0.58 |
| **Whole-Body** | 1 | 12.82 ± 3.28 | 2.15 ± 1.24 | 8.14 ± 2.35 | 1.66 ± 0.24 |
|  | 2 | 19.10 ± 2.25 | 1.85 ± 0.49 | 15.03 ± 1.31 | 1.70 ± 0.17 |
|  | 4 | 38.23 ± 8.80 | 5.47 ± 1.34 | 24.13 ± 2.80 | 4.58 ± 1.08 |
